# Vertical transmission of fungus-growing ant microbiota is species-specific and constrained by queens

**DOI:** 10.1101/2024.06.11.598432

**Authors:** Victoria A. Sadowski, Panagiotis Sapountzis, Pepijn W. Kooij, Jacobus J. Boomsma, Rachelle M.M. Adams

## Abstract

Multipartite symbioses are inherently complex, involving dynamic ecological interactions between organisms with intertwined yet distinct evolutionary histories. The fungus-growing (attine) ants facilitate maintenance of a symbiotic species network through maternal vertical transmission of an obligate fungal symbiont. While the gut microbiomes of fungus-growing ant species are remarkably simple, their fungal gardens support diverse microbial communities. Here, we focus on an understudied transmission bottleneck: the fungal garden pellet that nest-founding queens transfer to inoculate a new fungal garden. We used 16S rRNA metagenomic sequencing to reconstruct the extent of vertical transmission of bacteria to new gardens via queen pellets in four sympatric fungus-growing ant species (*Atta sexdens*, *Atta cephalotes*, *Acromyrmex echinatior*, and *Mycetomoellerius mikromelanos*) from Central Panama. We also characterized the bacterial communities associated with queen eggs and somatic tissues (mesosomas, guts and ovaries) to assess whether queens are likely to transmit symbiotic bacteria of workers, such as cuticular Actinobacteria and endosymbionts (*Wolbachia*, *Mesoplasma*, and *Spiroplasma*). Our results suggest that garden-associated bacteria are mainly horizontally acquired as the bacterial communities of pellets shared few bacterial taxa with the mature gardens of the four ant species investigated. While the bacterial communities of garden pellets showed some species-specificity, a subset of prevalent bacterial taxa were shared across ant species. Further, our findings provide evidence for vertical transmission of species-specific endosymbiotic bacteria through a transovarial route and/or via fecal droplets. Overall, while we found mixed evidence for vertical transmission of garden bacteria, our results support maternal transmission as a primary route for gut-associated symbionts. While our results suggest that vertical transmission of fungus-growing ant bacterial associates is mediated by the ant hosts, the mechanism behind this host control is not yet understood.

## Introduction

A fundamental question in the study of symbioses is how mutualistic partners associate in manners that allow mutual co-adaptation (1–3). Theory predicts that faithful vertical transmission will always allow positive selective feedback, yet there are many examples of horizontal transmission permitting reliable partner association (4–8). Social insects often maintain mutualistic relationships through both vertical transmission and repeated exchanges among nestmates, which is a form of social vertical transmission (9,10). The fungus-growing attine ants (Hymenoptera: Formicidae: Attini: Attina) maintain an obligate nutritional mutualism with mainly *Leucoagaricus* fungal cultivars (Basidiomycota: Agaricomycetes: Agaricales: Agaricaceae) reared in underground gardens and provisioned with plant material (11–13). The fungal garden functions as an external digestive system that degrades the plant material, with the help of a diverse assemblage of microbes. The carbon-rich organic material then produces accessible nutrients for the ants (14–19).

Dispersing gynes (female reproductives destined to become queens) provide a vertical transmission route for microbial symbionts in this symbiosis. During their mating flight, they use a pocket (infrabuccal pouch) in their oral cavity to vector fungal cultivar inoculum and associated microbes (i.e., a so-called infrabuccal pellet) from the natal nest to a new incipient colony (11). The gardens of *Atta* and *Acromyrmex* leafcutter ant species are known to host similar bacterial taxa (20), suggesting that these microbes are vertically transmitted via the pellet or the queen gut. However, direct evidence of vertical transmission of garden bacteria is lacking and horizontal acquisition from the environment cannot be excluded. How gynes obtain their pellet is unknown, but they are likely under positive selection to transmit beneficial garden bacteria that promote nutritional efficiency and to avoid transmission of pathogens or contaminants (but see Leftwich et al. (2020)). Dispersing gynes might preferentially select a pellet inoculum with a high abundance of bacterial symbionts. Alternately they may randomly acquire maternal fungus garden. Pellet surveys so far have suggested that both the fungal cultivar and the associated bacteria may indeed be vertically transmitted. For example, the infrabuccal pellets in newly mated queens of the leafcutter genus *Atta* contain bacteria that are closely related to bacteria commonly found in mature gardens (17,21,22). However, no studies have sampled pellets from more basal attine ants in which garden-associated bacterial communities are largely unknown (23–25).

Characterization of fungus-growing ant worker microbiomes has mostly focused on cuticular Actinobacteria which grow on worker mesosomas and produce antifungal compounds effective against *Escovopsis* fungal parasites and other pathogens (26–30). Additionally, the presence of visible cuticular Actinobacteria in fungus-growing ant species correlates with gut microbiota composition (Sapountzis et al., 2019). In the phylogenetic crown group, also referred to as the higher attine ants, worker gut bacterial communities have very low diversity compared to fungal gardens, similar to other insects with specialized diets (Colman et al., 2012; Yun et al., 2014; Sapountzis et al., 2019), and are usually dominated by a few OTUs belonging to *Wolbachia* and the bacterial orders Rhizobiales and Entomoplasmatales (Sapountzis et al., 2015, 2019; Zhukova et al., 2017, 2022).

These abundant bacteria persist when ant colonies are transferred from the field to the lab, suggesting they are functionally important symbionts (Sapountzis et al., 2015, 2019). *Wolbachia*’s role remains elusive although these bacteria are consistently found in somatic tissues of *Acromyrmex* species (Andersen et al., 2012; Frost et al., 2010). *Mesoplasma* (Mycoplasmatota: Mollicutes: Entomoplasmatales: Entomoplasmataceae) is primarily associated with leafcutter ant species while non-leafcutter attines also associate with *Spiroplasma* (Mycoplasmatota: Mollicutes: Entomoplasmatales: Spiroplasmaceae) (31,34,37). These Entomoplasmatales symbionts likely provide multiple metabolic services to their attine host ants by converting citrate into easily accessible acetate and by recycling excess—fungal cultivar produced—arginine into ammonia (38,39). Outside of these primary endosymbionts, Rhizobiales often form biofilms in the guts of leafcutter ants, possibly contributing to mutualism-stability by nutrient exchange with other gut symbionts (34,40). However, it remains unclear whether these bacterial symbionts are vertically transmitted by queens. Previous work has shown that in *Acromyrmex, Wolbachia* is transmitted via eggs, but Entomoplasmatales (*Mesoplasma* and *Spiroplasma*) symbionts were instead primarily transmitted from older to newly emerged workers (10). Entomoplasmatales have also been detected in the eggs of the non-leafcutter attine *Sericomyrmex amabilis* and in the gynes of its social parasite (37), indicating the potential for a mixed mode of transmission, vertically from parent to offspring and horizontally between ant species. Thus, while recent studies have begun to establish the functionality of these symbioses in attine ant workers, further investigation is needed to better understand their transmission dynamics.

In this study we used 16S ribosomal RNA metabarcoding sequencing to characterize bacterial microbiota in four fungus-growing ant species from the Panama Canal region: *Atta sexdens* (Linnaeus, 1758) (41), *Atta cephalotes* (Linnaeus, 1758) (41), *Acromyrmex echinatior* (Forel, 1899) (42) and *Mycetomoellerius mikromelanos* Cardenas, Schultz, & Adams, 2021(43). We sampled gyne tissue (used as a proxy for queens), pellets, and garden from mature colonies in all four species. We also sampled tissue from young newly mated queens, garden and eggs in *A. sexdens* and *M. mikromelanos* colonies. In our hypotheses we use “queens” to refer to gynes and queens. In the following sections, we specify either gyne or queen samples where appropriate.

First, we made within-species comparisons to test the hypothesis (H1) that attine queens vertically transmit garden bacteria through their pellet inocula. Under this hypothesis we expected that fungal pellet bacterial communities would consistently have the same bacterial taxa found in fungal gardens. Alternatively, but not mutually exclusive, we also hypothesized (H2) that ant-associated bacteria are vertically transmitted by the queens themselves. Under this hypothesis we expected that known gut- and endosymbionts (*Wolbachia, Mesoplasma* and *Spiroplasma*) and cuticular Actinobacteria (in *Acromyrmex* and *Mycetomoellerius*) would be found in tissues of dispersing and founding queens. We further used a comparative framework to explore bacterial community diversity between the four sympatric ant species. Here, we hypothesized (H3) that vertically transmitted microbiota should have signatures reflecting host taxonomy. Under this hypothesis, we expected that queens of the phylogenetically basal (non-leafcutter) *M. mikromelanos* would host unique bacteria relative to the three more closely related leafcutter ant species (13).

## Results

### Alpha-diversity and community composition within species

Analysis of rarefaction curves at 97% sequence similarity indicated that sequencing coverage was sufficient, apart from a small number of samples with exceptionally high diversity (S1 Fig). Alpha-diversity, or within-sample species richness and diversity, was assessed using the Inverse Simpson index, which showed that diversity was variable across samples within ant species, with gardens and pellets not necessarily being more diverse than gyne or queen associated tissues (ovaries, gut, and mesosoma) (S2 Fig, S5 Table). The composition of bacterial phyla was also variable across sample types, although all bacterial communities were composed mostly of Pseudomonadota, Mycoplasmatota, Actinomycetota and Bacteroidota (S3 Fig). Pseudomonadota and Mycoplasmatota were particularly dominant in the bacterial communities of *Acromyrmex echinatior* (S3 Fig), which was largely due to the high abundance of *Wolbachia* and Entomoplasmatales symbionts in this species.

### Vertical transmission of garden-derived bacteria

Using beta-diversity analyses to explore differences between types of samples, we found that pellets and gardens hosted distinct bacterial communities in all four ant species (Fig 1, S6 Table, S7 Table). This suggests that mature gardens obtain additional bacteria not present in the pellet inocula, therefore, our H1 hypothesis that newly mated queens vertically transmit garden bacteria was not strongly supported. This interpretation became apparent from the few OTUs that were core taxa (i.e., present in >65% of samples) in both pellet and garden communities (S4 Table). In *Atta cephalotes*, *Mesoplasma* (OTU2), Rhizobiaceae (OTU8), and *Pelomonas* (OTU21) were shared core taxa although there were no such shared core OTUs in *Atta sexdens* pellet and gardens (S4 Table). The Rhizobiaceae (OTU8) was 100% identical to an earlier found 16S rRNA sequence from gut bacteria of South American *Atta laevigata* (GenBank accession: KF250158.1) and to bacteria shared with other fungus-growing ants (*Sericomyrmex amabilis* and *Mycetomoellerius zeteki*) and their social parasites (*Megalomyrmex symmetochus* and *Megalomyrmex adamsae* respectively) (GenBank accession: LC027777.1; Liberti et al. (2015)). A previously established garden symbiont, *Klebsiella* (OTU6) was a core taxon in both pellet and garden communities in both *Ac. echinatior* and *M. mikromelanos* (S4 Table), but pellets and gardens remained distinct in beta-diversity measures due to relative abundance of other OTUs (Fig 1C, D). Surprisingly, some known ant-associates (*Mesoplasma*, *Wolbachia* and *Actinobacteria*) were also core taxa of pellet and gardens for all four species (S4 Table). Thus, some ant endosymbionts may be transmitted via pellets along with a few garden-associated bacteria.

**Fig 1.**
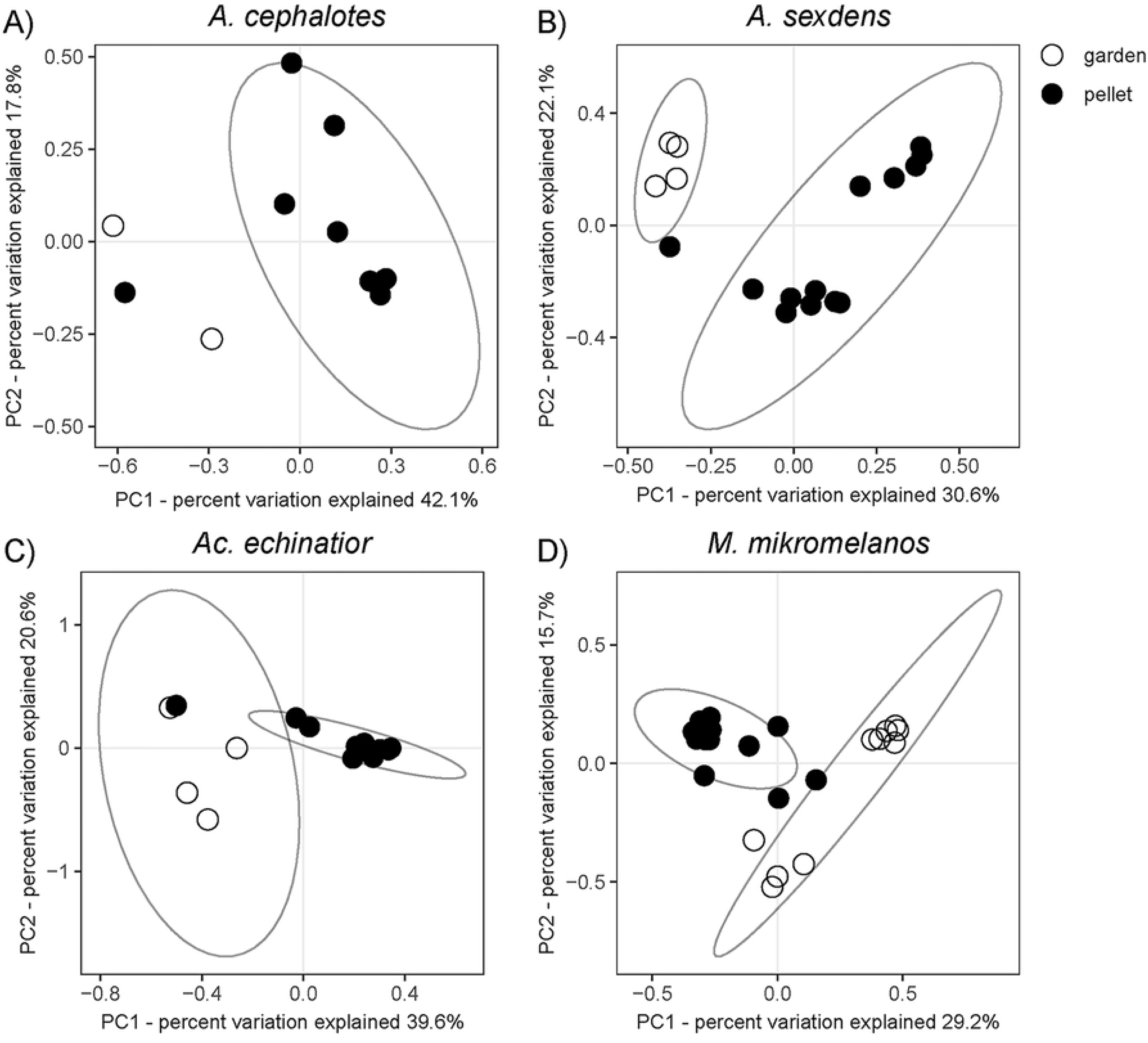
Within-species comparisons of pellet and garden beta-diversity using Weighted Unifrac Principal Coordinate Analysis (PCoA). (A) *Atta cephalotes* (B) *Atta sexdens*, (C) *Acromyrmex echinatior* and (D) *Mycetomoellerius mikromelanos.* Ellipses represent 95% confidence intervals. The sample size for *A. cephalotes* garden samples was too low to calculate a statistical ellipse, but pellets and gardens hosted different bacterial microbiomes in all four species (p<0.05).

### Mechanisms of vertical transmission of known ant symbionts

Our data supported our H2 hypothesis that ant-associated bacterial gut symbionts can be vertically transmitted by founding queens. For two species, we were able to sample young foundress queens in addition to unmated gynes from older colonies. *Atta sexdens* and *M. mikromelanos* queens hosted some of the same bacterial taxa in their ovaries as were consistently found in their eggs (S4 Table). Beta-diversity, egg and ovary bacterial communities were somewhat distinct (Fig 2, S6 Table, S7 Table). However, these differences in beta-diversity between eggs and ovaries were driven primarily by lower relative abundance of the known symbionts in the eggs as compared to the ovaries, so the overall consistency of vertical transmission appears to be a more important result than the slight differences. In order to preserve queen-derived microbes coating the eggs, they were not surface sterilized before DNA extraction. Thus, contamination with garden-derived microbes could drive differences between the microbiota of eggs and ovaries as well.

**Fig 2.**
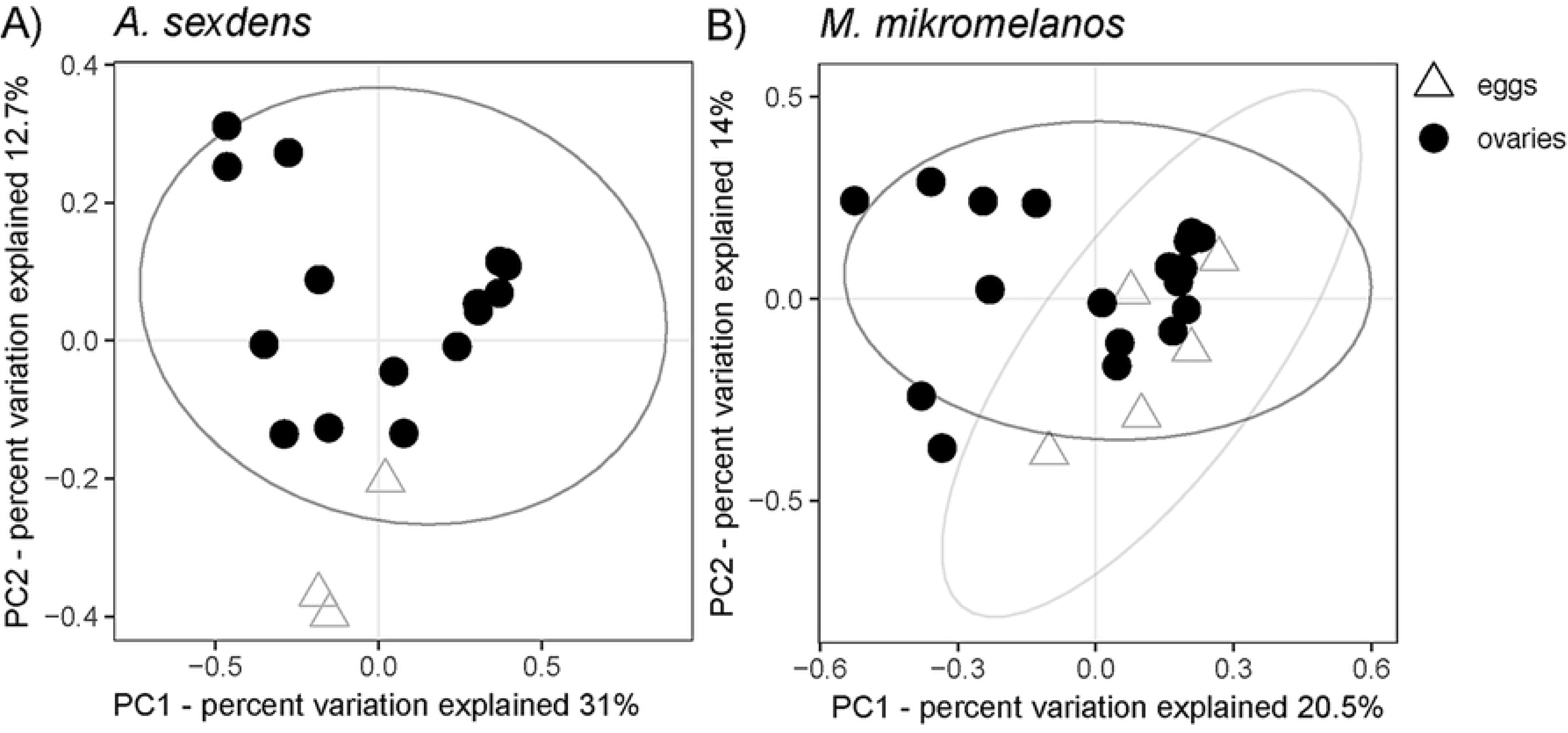
Within-species comparisons of egg and ovary beta-diversity using Weighted Unifrac Principal Coordinate Analysis (PCoA). (A) *Atta sexdens* and (B) *Mycetomoellerius mikromelanos.* Ellipses represent 95% confidence intervals. The sample size for *A. sexdens* eggs was too low to calculate a statistical ellipse, but eggs and ovaries hosted overlapping and statistically different bacterial microbiomes in both species (p<0.05).

Known bacterial endosymbionts were consistently found in/on gyne and queen tissues and were represented by the three most abundant OTUs in our dataset, belonging to the genera *Wolbachia*, *Mesoplasma* and *Spiroplasma* (Fig 3). While we recovered a number of rare additional OTUs from the same bacterial genera, a single OTU was always abundant for each symbiont genus. The most abundant OTU across all samples, *Wolbachia*, was part of the core community in all queen- or gyne-associated tissues of *A. sexdens* and *Ac. echinatior*, but not *A. cephalotes* (S4 Table), and was abundantly found in some samples (Fig 3). The second most abundant OTU across all samples was *Mesoplasma* which shares 99% sequence similarity with a previously identified symbiont of fungus-growing ants, *EntAcro1* (GenBank accession KR336618) (Sapountzis et al. 2015). *Mesoplasma* constituted a large proportion of the bacterial communities associated with *A. cephalotes* and *Ac. echinatior* queen guts and ovaries (Fig 3).

**Fig 3.**
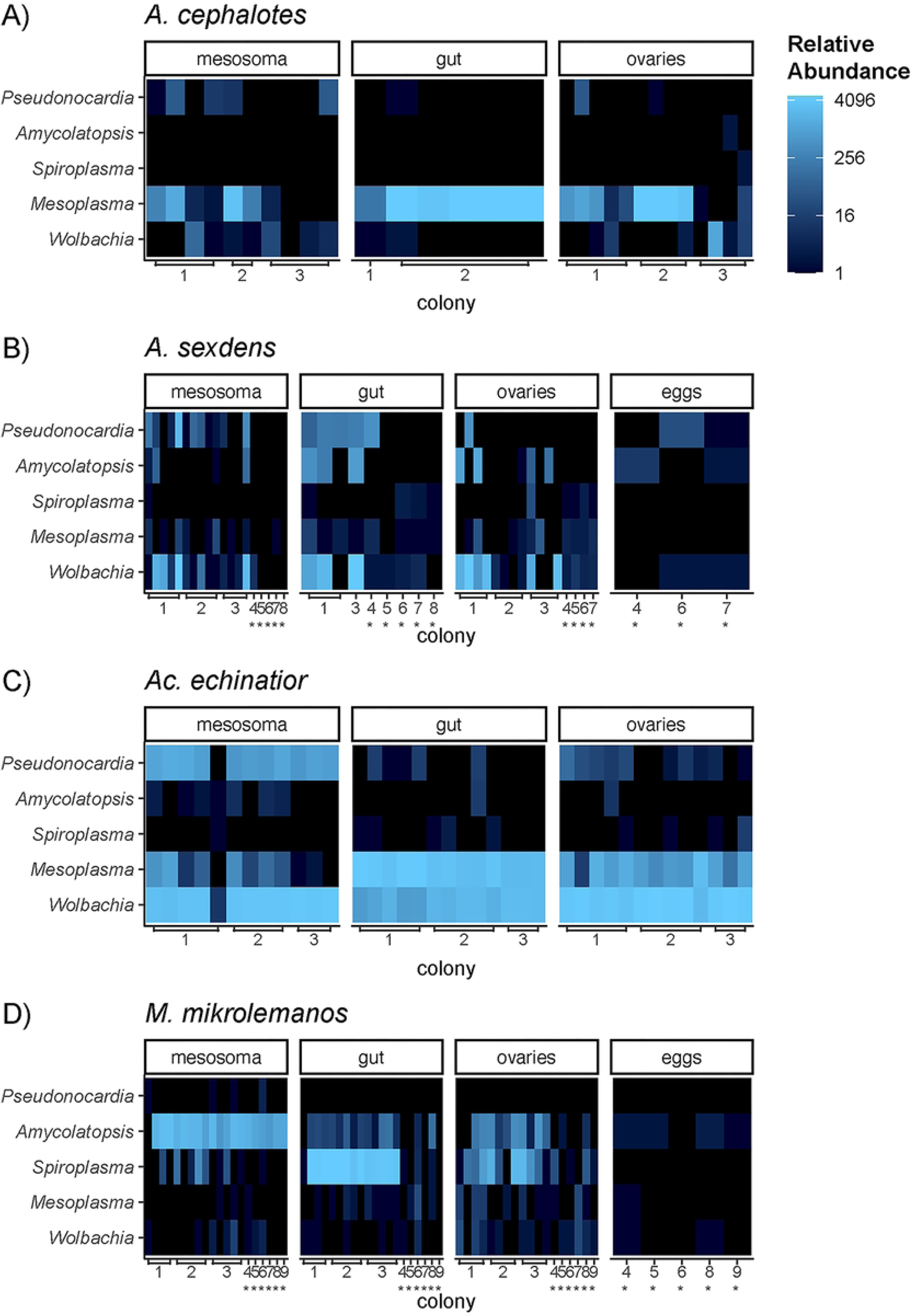
Heatmaps depicting interspecific variation in bacterial symbiont abundance among queen tissues. (A) *Atta cephalotes* (B) *Atta sexdens*, (C) *Acromyrmex echinatior* and (D) *Mycetomoellerius mikromelanos.* Each row represents a single OTU (*Pseudonocardia* OTU9, *Amycolatopsis* OTU5, *Spiroplasma* OTU3, *Mesoplasma* OTU2 and *Wolbachia* OTU1) and each column represents an individual queen or gyne ant. Samples from young queens, rather than gynes, are noted with asterisks (*). The color scale is based on log base 4 transformation of relative (proportional) abundance out of 5000 reads. Colony names are simplified as numbers. See S1 Table for corresponding colony collection codes.

This OTU was a core taxon in queen and gyne guts for all three leafcutter species (S4 Table) but was less abundant in *A. sexdens*. Although the same *Mesoplasma* OTU was sometimes identified in *M. mikromelanos*, *Spiroplasma* (OTU3) was more consistently present and a member of the core community in queen and gyne guts and ovaries of this species (Fig 3D). This OTU shared a high sequence similarity with the *Spiroplasma* in the guts of *M. zeteki* and *S. amabilis* workers (∼98%; Liberti et al. 2015). In *A. sexdens*, queen ovaries and their eggs shared a number of core bacterial taxa including *Wolbachia*, but neither *Spiroplasma* nor *Mesoplasma* were shared core taxa in the ovaries and eggs. The latter also applied in the case of *M. mikromelanos* (S4 Table). However, consistent bacterial symbionts (*Wolbachia* and *Spiroplasma*) in the ovaries of *A. sexdens* and *M. mikromelanos* were present in low relative abundances in the egg samples of the same queens (Fig 3).

Actinobacteria were present in queen and gyne mesosoma samples of all ant species but were most abundant in *M. mikromelanos* and *Ac. echinatior* (Fig 3; S3 Fig). The primary genus of Actinobacteria associated with fungus-growing ants, *Pseudonocardia* (OTU10), was a core member of the *Ac. echinatior* queen mesosoma microbiome (S4 Table) and was highly abundant in that species (Fig 3C). A close relative to *Pseudonocardia* and another symbiont of fungus-growing ants, *Amycolatopsis* (OTU5), was a core member of all queen and gyne tissues in *M. mikromelanos* (S4 Table) and was consistently abundant in the mesosoma samples of this species (Fig 3D). Actinobacteria made up a smaller proportion of the overall mesosoma community in *Atta* queens/gynes and were not among the core taxa in those species (S3 Fig, S4 Table), yet the same two Actinobacterial OTUs mentioned above were components of queen/gyne bacterial microbiota in every *Atta* colony sampled and were sometimes relatively abundant (Fig 3). The *Pseudonocardia* (OTU9) was >97% similar in sequence to *Pseudonocardia* previously isolated from *Trachymyrmex* (44), *Acromyrmex* and *Apterostigma* (45).

### Pellet bacterial communities have the same genera across species yet maintain their host-ant-specific signatures

While hypotheses H1 & H2 address vertical transmission of garden and ant-associated symbionts, respectively, H3 emphasizes possible interspecific similarities and differences in vertically transmitted bacterial communities. Five OTUs were identified as core members of pellet communities in all four ant species sampled: *Moraxella* (OTU14), *Staphylococcus* (OTU15), *Lawsonella* (OTU20), *Pelomonas* (OTU21), and *Micrococcus* (OTU27) (S4 Table). None of these five have been previously identified as fungus-growing ant symbionts, but *Staphylococcus* has been found in earlier fungal garden studies (22,23,46). We found no significant pairwise differences in garden microbiota between species aside from between *M. mikromelanos* and *Ac. echinatior* (S6 Table, S7 Table), but across all four species we found that the composition of pellet bacterial communities differed significantly (S6 Table; S7 Table).

Specifically, post-hoc tests show that *M. mikromelanos* pellet bacterial communities were significantly different from those of the three leafcutter species (p=0.006), and that pellets of *Atta* species were more similar to each other than any pairwise comparison between different genera (Fig 4, S6 Table, S7 Table). However, pellets from two colonies of *A. sexdens* shared similar bacterial microbiota with *Ac. echinatior*, so the *A. sexdens* samples appeared as two distinct clusters in the PCoA plot (Fig 4). Pellet bacterial communities were not significantly different between colonies within each ant species, apart from the above-mentioned two colonies of *A. sexdens*. Additionally, the other colony of *A. sexdens* (As1) had pellet communities more similar to those of *A. cephalotes* (Fig 5).

**Fig 4.**
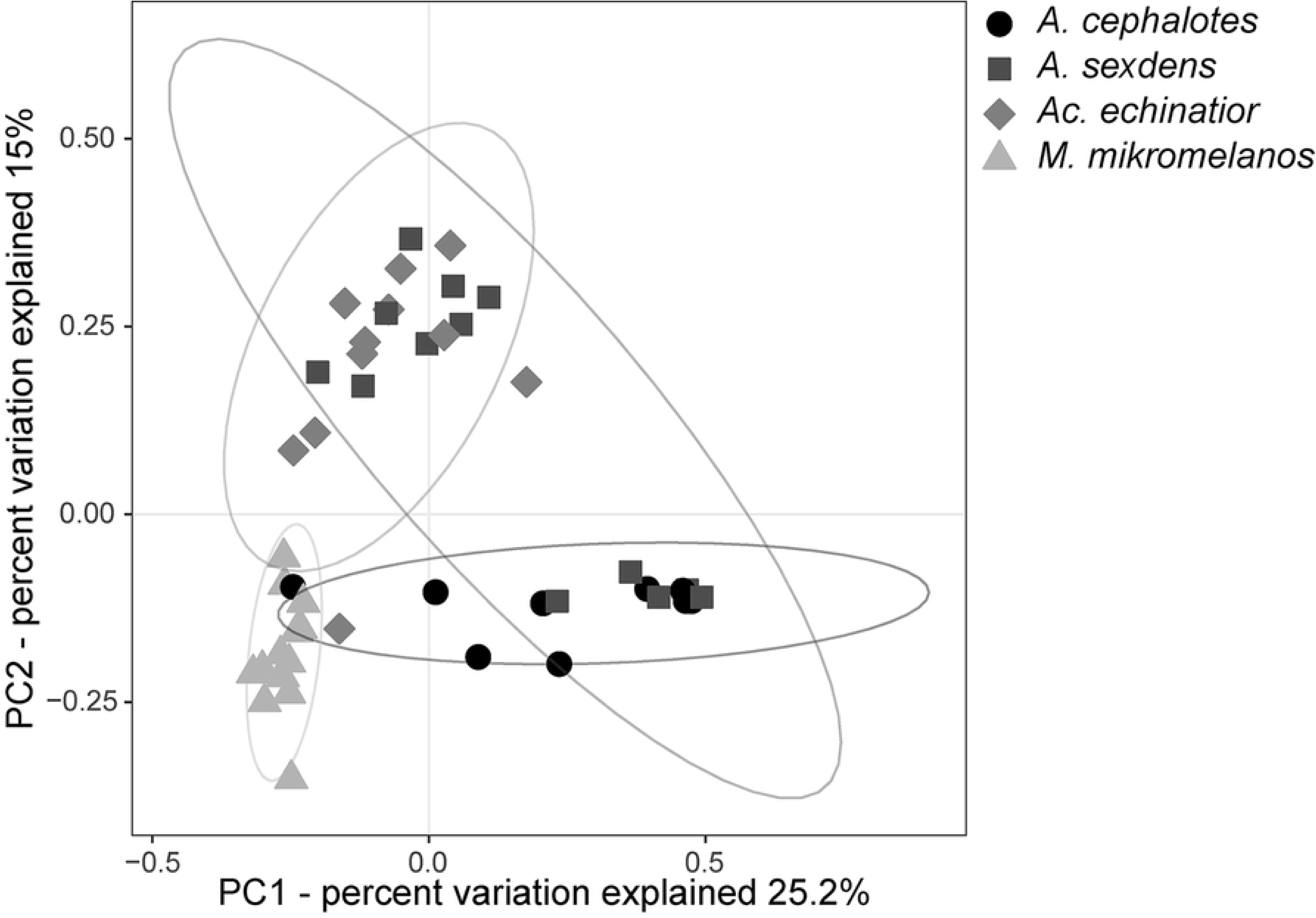
Between-species comparisons of pellet beta-diversity using Weighted Unifrac Principal Coordinate Analysis (PCoA). Shapes and shades of black/grey designate the ant species from which individual pellets were sampled (*Atta cephalotes*, *Atta sexdens*, *Acromyrmex echinatior*, and *Mycetomoellerius mikromelanos*) and ellipses represent 95% confidence intervals. *M. mikromelanos* pellets hosted different bacterial microbiota than those of the three leafcutter species (p=0.006).

**Fig 5.**
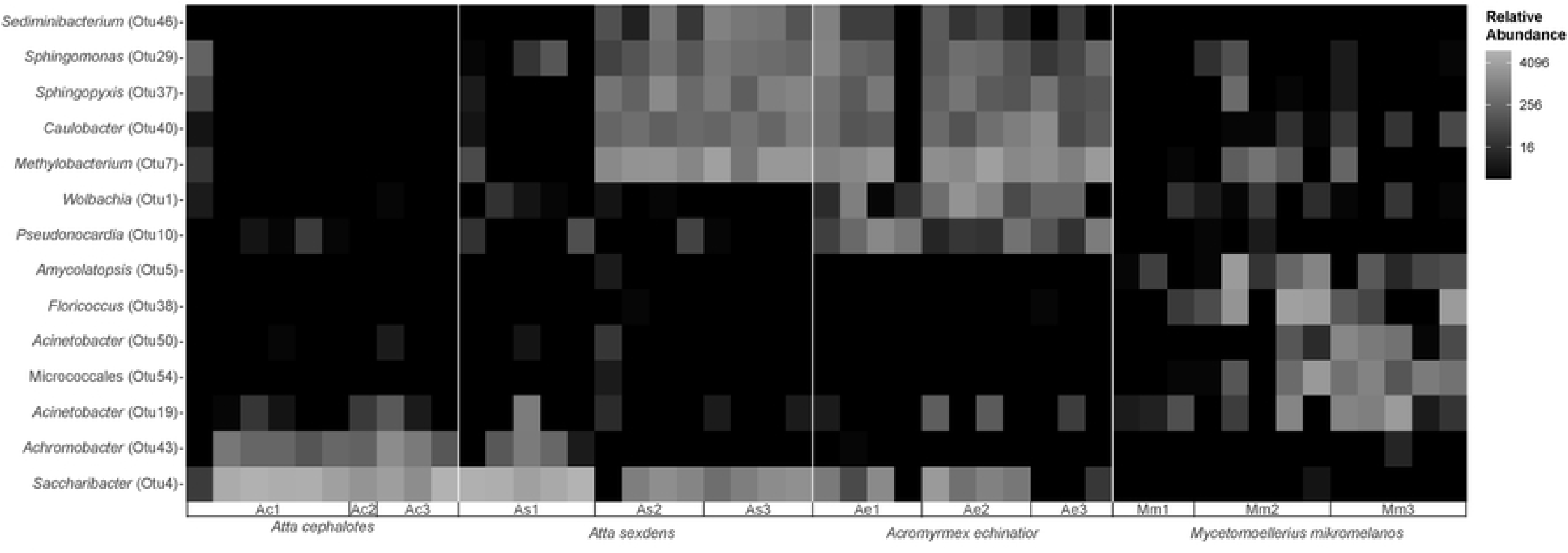
Heatmap depicting variation in abundance of bacterial OTUs driving interspecific differences in pellet microbiota. Only the 14 OTUs which contributed significantly to differences in beta-diversity between species are shown. The Y-axis is ordered by metric multidimensional scaling (MDS) using weighted Unifrac distances. The color scale is based on log base 4 transformation of relative (proportional) abundance out of 5000 reads. See S1 Table for corresponding colony collection codes.

SIMPER analysis captured that the differences in beta-diversity between *M. mikromelanos* pellet bacterial communities and those of the other three ant species were driven by a few abundant bacterial taxa, primarily *Saccharibacter* (OTU4), *Amycolatopsis* (OTU5), *Floricoccus* (OTU38) and an unidentified Micrococcales (OTU54) (Fig 5; S8 Table).

Differences between *Ac. echinatior* pellet communities and those of *A. cephalotes* were driven primarily by the relative abundance of *Saccharibacter* (OTU4) and *Methylobacterium* (OTU7) (Fig 4; S8 Table). Particularly OTU 4 (*Saccharibacter*) contributed significantly to differences in pellet beta-diversity across ant genera and accounted for 3.5-25% of dissimilarity between these communities (S8 Table). *Saccharibacter* (OTU4) is a member of the core pellet community for all three leafcutter species (S4 Table) and was particularly abundant in gyne pellets of some *Atta* colonies (Fig 5). OTU4 is most closely related to uncultured bacteria isolated previously from leafcutter ant bodies and refuse dumps (99.2% similarity to GenBank accessions KF248837.1 & LN568338.1) but relatively distantly related to any other GenBank reference sequences (∼91% sequence identity to *Saccharibacter floricola*; S2 Table). While *Saccharibacter* was not abundant in pellets of *M. mikromelanos*, the lactic acid bacterium *Floricoccus* (OTU34) was exclusively found in *M. mikromelanos* pellets and is closely related to bacteria obtained from plant isolates (47). Several other lactic acid bacteria were found to be generally abundant in our dataset: *Weissella* (OTU12), another *Floricoccus* (OTU11), and *Staphylococcus* (OTU50). OTU54 shared <94% sequence similarity with deposited GenBank sequences except for an uncultured bacterium from workers of an *Aphaenogaster* ant species (Russell et al., 2012), but this OTU could only be classified at the Order level (Micrococcales).

We also found host-specific signatures in microbial communities associated with queen and gyne ovaries, guts and mesosomas (Fig 6). While egg bacterial microbiota were similar between those of some *Atta* colonies and *M. mikromelanos*, the ovaries hosted distinctly different microbiota across all four ant species despite sharing some core taxa (e.g., *Mesoplasma* and *Wolbachia*) (Fig 6A; S6 Table, S7 Table). The beta-diversity analysis for queen and gyne mesosoma microbiota showed genus specific patterns, visualized by distinct clusters for *M. mikromelanos* and *Ac. echinatior* and a lack of differentiation between the two *Atta* species who do not consistently host Actinobacteria (Fig 6B; S6 Table, S7 Table). Also, gut microbiota were distinct between all four ant species (p<0.05) despite sharing of *Wolbachia* and *Mesoplasma* across the three leafcutter species (Fig 6C; S6 Table). Overall, we found that fungus-growing ant queen and gyne ovaries, guts and mesosomas hosted species and/or genus-specific bacterial communities even though some bacterial taxa were shared across species.

**Fig 6:**
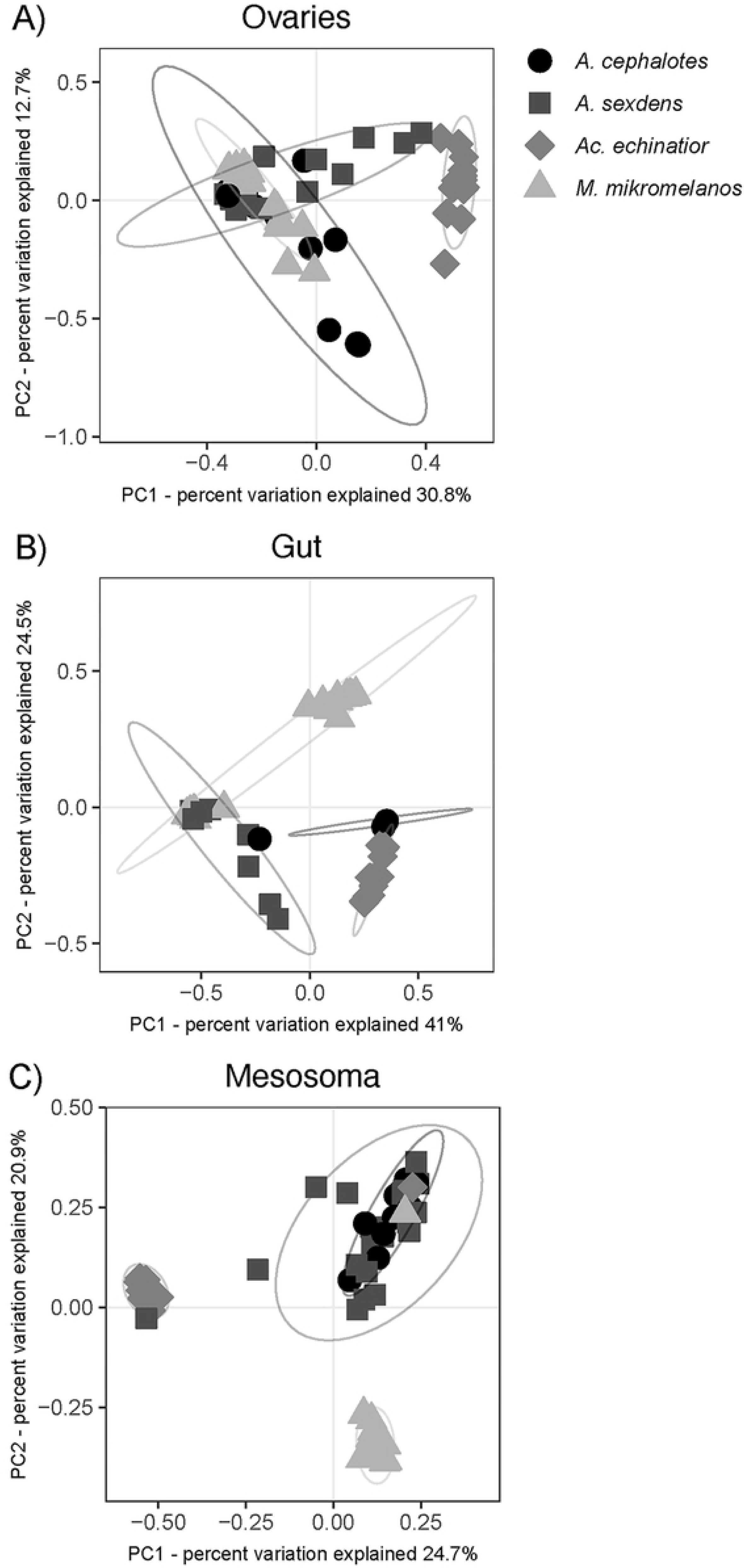
Between-species comparisons of beta-diversity across queen/gyne tissues using Weighted Unifrac Principal Coordinate Analysis (PCoA). (A) Gynes/queens of all species hosted distinct bacterial communities in their ovaries (p<0.05). (B) Gynes/queens of all species also hosted distinct bacterial communities in their guts (p<0.05). Four *A. cephalotes* gyne gut samples from colony Ac2 are directly overlaid in the plot as these samples were dominated by *Mesoplasma*. (C) The mesosomas of the two species of *Atta* were indistinguishable (p=1). *Ac. echinatior* gynes and *M. mikromelanos* gynes/queens hosted distinctly different bacterial communities on their mesosomas (p=0.006). Samples of gynes and queens from *A. sexdens* and *M. mikromelanos* are analyzed together. Ellipses represent 95% confidence intervals.

## Discussion

In this study we tested three hypotheses to develop a better understanding of the transmission dynamics of fungus-growing ant bacterial microbiota. Despite high bacterial diversity in ant gardens, our results indicate that only a select few bacteria are transmitted by dispersing and colony founding queens, supporting our first hypothesis (H1). In particular, *Klebsiella*—a nitrogen fixer in *Atta* and *Acromyrmex* ant gardens (21)—appears to be transmitted via queen pellets in *Ac. echinatior* and *M. mikromelanos.* The major ant-associated symbionts, *Wolbachia*, *Mesoplasma* and *Spiroplasma*, were also consistently found in pellet and garden samples, indicating that the fungal garden may serve as a reservoir for these symbionts and/or they may play a symbiotic role in the garden environment (22,49). Because fungus-growing ant queens deposit fecal droplets when feeding and tending their garden, both queen gut bacteria and pellet inocula likely contribute to the assembly of bacterial communities in incipient fungal gardens. Experimental inoculation with genetically modified bacterial strains that express visible markers (e.g., fluorescent proteins) could be used to confirm the inferred pellet-mediated vertical transmission of these bacteria to new gardens. Strain-level assays, such as culturomics using generic and species-specific media, coupled with single-cell metagenomics could also be used to unravel the hidden diversity of individual bacterial strains that are transmitted by founding queens from natal gardens.

Overall, our results support our second hypothesis that attine ant queens vertically transmit some OTUs corresponding to known bacterial symbionts that in previous studies have been identified as co-adapted mutualists of the farming symbiosis (21,31,36,39). In each species, we found that only one strain of *Wolbachia*, *Mesoplasma* and/or *Spiroplasma* was highly abundant and often shared with ant species of the same or a sister genus. Other bacteria previously associated with fungus-growing ants, such as Rhizobiales, were not consistently found. Our results corroborate previous work suggesting complete transovarial transmission of *Wolbachia* but not of Entomoplasmatales (e.g, *Mesoplasma* and *Spiroplasma*) in the phylogenetic attine crown group of the leafcutter ants (10). However, contrary to a recent study, which suggested that *Atta sexdens* from Brazil do not consistently associate with *Mesoplasma*, we found these bacteria to be a core taxon in the guts of *A. sexdens* and *A. cephalotes* queens and/or gynes (50). While Entomoplasmatales symbionts may not be transmitted consistently via eggs, their abundance and prevalence in gardens, worker guts, and queen and gyne guts remains highly relevant. We hypothesize that trophallaxis between adults and reproductive larvae mediates the transmission of Entomoplasmatales symbionts, as shown for transmission of bacteria in other social arthropods (9,51,52).

We also found that *Acromyrmex* and *Mycetomoellerius* queens vertically transmit Actinobacteria which are maintained as biofilms through glandular secretions on the exoskeleton (Andersen et al., 2013, 2015; Barke et al., 2011; Sapountzis et al., 2019). The same Pseudonocardiaceae (Actinobacteria: Actinomycetales) OTUs were found across all four ant species, despite the lack of actinobacterial biofilm on *Atta* cuticles. Interestingly, recent evidence suggests that cuticular microbiota in *Atta* do not play a defensive role, at least against the *Metarhizium anisopliae* fungal entomopathogen (53). While multiple Actinobacterial strains can occur on fungus-growing ant worker cuticles, we found that only one or two strains dominated the bacterial communities of queen and gyne mesosomas, similar to what has been previously shown on cuticles of callow workers of *Acromyrmex* leafcutter ants (28). It is important to note that *Pseudonocardia* has also been found in the guts of sympatric Panamanian *M. zeteki* (31), so results obtained from whole (parts of) body extracts should be interpreted with caution.

Despite ample opportunities for fungal gardens (and thus infrabuccal pellets) to acquire bacterial associates horizontally from the environment, we identified a set of core pellet bacterial OTUs with as of yet unknown functions that were shared across the four Panamanian fungus-growing ant species that we studied (*Moraxella*, *Staphylococcus*, *Lawsonella*, *Pelomonas*, and *Micrococcus*). *Pelomonas* (OTU 21) is closely related to bacterial OTUs from a variety of sources, but particularly relevant in the context of our study is the association of this bacterial genus with fungal gardens of *Mycocepurus goeldii* (54) and the males of *Trachymyrmex septentrionalis* (25). *Staphylococcus* may function to increase bioavailable iron and promote growth of the fungal cultivar (46). In general, these bacterial genera are closely related to human-associated or environmentally abundant isolates.

Notably, one colony of *A. sexdens* (As1) shares several OTUs with *A. cephalotes* colonies whereas the other two colonies, As2 and As3, share different OTUs with *Ac. echinatior*. All three species are leafcutters and small fragments of plant material are visible in their pellets, thus these similarities may be influenced by the bacteria associated with the plants the ants are gathering to feed their cultivar. Still, the consistency of these as well as *Moraxella*, *Staphylococcus*, *Lawsonella*, *Pelomonas*, and *Micrococcus* bacterial taxa in infrabuccal pellets of attine dispersing queens suggests they could be facultative symbionts but this remains to be tested. We believe the presence of these bacteria is unlikely to be a result of contamination as they’re not found in our sequencing blanks nor consistently in other sample types. Further, this shared set of core bacterial taxa in pellets across attines suggests that there may be selective factors conserved across attines that influence which garden bacteria are vertically transmitted. In this context it seems striking that we found so little evidence for garden-associated symbionts of known function to be vertically transmitted. Overall, our results corroborate previous evidence that bacterial associates of fungal gardens are acquired through both vertical and horizontal transmission (55).

Although we identified a shared set of core bacteria across the infrabuccal pellets of all four species investigated, we also found that the bacterial microbiota of *M. mikromelanos* pellets were distinct from those of the three leafcutter ants, supporting our third hypothesis (H3). This specificity in pellet microbiota mirrors the divergence in attine ant abdominal microbiomes that arose when the ancestors of the *Paratrachymyrmex* sister genus of *Mycetomoellerius* colonized Central and North America (Branstetter et al. 2017; Sapountzis et al. 2019). The significant ecological differences between *M. mikromelanos* and sympatric leafcutter ants may also have influenced the vertical transmission dynamics of their microbiota. For instance, fungal cultivar clade, ant microbiomes, and preferred forage material are systematically different between *M. mikromelanos* and the three leafcutter ant species of this study (12,31,57) The ants included in our study cultivate fungi from two sister clades, with the three leafcutter species cultivating a single species, i.e., *Leucoagaricus gongylophorus*, and *M. mikromelanos* cultivating a related but undescribed species of fungus (13,58,59). There is some evidence that the relative abundance of bacterial symbionts is dependent on cultivar clade, or vice versa (22,49). The differences in pellet microbiota may thus be a consequence of the selective environment imposed by the fungal cultivar strain. While we did not find a strong ant host signature in the garden microbiota, our garden sample size was relatively small, so future work should include more within-nest samples considering the potential for undetected variation in bacterial communities (23).

## Conclusions

Overall, our results provide further evidence that the role of attine ant queen infrabuccal pellets is more complex than just vectoring an inoculate of a queen’s maternal fungal cultivar to the next generation. Recent studies have characterized the bacterial microbiomes of the digestive systems of attine ant workers (10,31,36) and fungal gardens of mature colonies (15,20,55,60,61). These have clarified that a number of bacterial strains have specific functions in the farming symbiosis, suggesting a close ecological relationship with their host ants (21,34,39,40). Such processes of mutual co-adaptation are easy to conceptualize when these bacteria are vertically transmitted, but much harder to explain when they are are acquired *de novo* from ecologically different attine ant habitats.

Our study is the first to target these questions by direct sampling of the eggs and infrabuccal fungus-garden pellets of multiple species of attine ants. While we sampled only four sympatric attine ant species from a single Panamanian location and our sample sizes were modest, the overall trend of our results is that bacterial symbionts with known complementary functions in this complex farming symbiosis are indeed vertically transmitted via queens through multiple routes. Although, some horizontal transmission of symbionts is likely, especially in fungal gardens. We also identified several bacterial lineages with unknown functions that are consistently vertically transmitted and where further work might elucidate a functional role. As this study serves as a proof-of-concept, future work can build from our comparative approach and test the generality of our results across a larger assembly of fungus-growing ants with a wider geographic distribution.

## Materials and Methods

### Sample collection

*Atta sexdens, A. cephalotes, Ac. echinatior*, and *M. mikromelanos* samples were collected from field nests in the Panama Canal region in May 2014. Gynes (unmated female reproductives) were collected from live colonies immediately or < 7 days after they were excavated from the field after which pellets were extracted from their oral cavity using sterilized forceps and preserved dry at −80°C until DNA extraction. Nest-founding queens were collected for *A. sexdens* (n=5) and *M. mikromelanos* (n=6) in the same time period. In these cases, eggs and incipient fungal garden material were sampled. These small gardens presumably originated from pellets that queens disperse with. Garden samples were collected from a queen-dug cavity and were fed and nurtured by the young queen. Some may have had plant material recently incorporated into the garden substrate. Fungal garden samples were collected directly upon field sampling and stored at −80°C until DNA extraction. Single guts (midgut and hindgut), and reproductive tracts for queens, were dissected from the rest of the body (i.e., mesosoma) in sterile phosphate-buffered saline (PBS) under a stereomicroscope to consider these microbiomes separately. Although our target in dissecting queen reproductive tracts was the ovaries, connected reproductive tract organs (i.e., spermathecae) were not removed to preserve sample integrity.

### Molecular methods

DNA was extracted from frozen guts, ovaries, mesosomas, and eggs using the DNeasy blood and tissue kit (Qiagen) following the manufacturer instructions. DNA from frozen fungal garden and pellet samples was extracted with a Cetyltrimethylammonium Bromide (CTAB) solution followed by a phenol–chloroform extraction (62). One negative control (i.e., MilliQ water) extraction was performed for each set of 28 samples-amounting to a total of 5 blanks. We also included a mock community (BEI Resources, Manassas, VA, USA) extraction to assess error rates. PCR primers 515F (5’-GTGCCAGCMGCCGCGGTAA −3’ and 806R (5’-GGACTACHVHHHTWTCTAAT −3’) covering the V4 region of the 16S rRNA gene were used for PCR reactions using a mix containing 2.0µl 10× AccuPrime™ PCR Buffer II (15 mM MgCl2), 0.15µl AccuPrime™ Taq DNA Polymerase (2 units/µl, Life Technologies), 1.0µl of each primer (10 µM), 1.0µl diluted template and autoclaved MilliQ water to a total of 20µl. The following thermal profile was used: 95°C for 2 min, followed by 30 cycles of 95°C for 20 s, 55°C for 15 s, 72°C for 5 min, and final extension at 72°C for 5 min. All samples were re-eluted in 50-150µl AE buffer (Qiagen) then quantified using an ND-1000 spectrophotometer (NanoDrop Technologies, Wilmington, DE, USA). The purified amplicons were then pooled in equimolar concentrations. Extracted DNA was sent to Microbial Systems Laboratory at the University of Michigan (Ann Arbor, MI) for library preparation and 16S Miseq Illumina sequencing as described in (63).

### Sequencing data processing

Sequences were processed with mothur v1.44.3 (64). The raw sequences were filtered using a minimum quality score of 25 and reads were discarded when longer than 275 bp in length or showing any ambiguous bases. Unique sequences were then aligned to a non-redundant database (SILVA v138) customized for the 16S rRNA V4 region and sequences were removed which did not cover the V4 hypervariable region or had a homopolymer length above 8. Unique sequences were screened again after removing overhangs. Chimeric sequences were identified using vsearch v. 2.13.3 and removed. Filtered sequences were identified taxonomically using the same customized SILVA database (with an 80% bootstrap confidence threshold) and any sequences classified as Archaea, Eukaryota, mitochondria, chloroplast or unknown were excluded. Bacterial sequences were clustered *de novo* into operational taxonomic units (OTUs) using the average neighbor algorithm with a 97% similarity threshold while retaining singletons. The final sequence file consisted of 9,271 filtered and unique OTUs. The taxonomy of the 50 most abundant OTUs was verified manually using NCBI BLASTn (S2 Table).

### Data analyses

Rarefaction and diversity analyses were completed in R v. 3.6.2 with the packages phyloseq v1.38 and vegan v2.5.7 using the OTU table, taxonomy and tree files imported from mothur (65,66). Rarefaction curves were visualized using vegan and ranacapa v1.0 R packages (67). Alpha-diversity, or within sample diversity, was evaluated using the Inverse Simpson index since the Shannon index is affected by low sequencing coverage.

Significant differences between groups were determined by non-parametric Kruskal-Wallis and *post hoc* Wilcoxon rank sum tests. All samples were subsampled to 5,000 reads, reducing the sample number to 268 (28 samples had <5000 reads) and reducing the total number of OTUs from 9,271 to 6,306. This dataset was used for all downstream analyses. Following the methods of Meirelles et al. (2016), we considered a bacterial OTU as part of the core microbiota if this taxon was recovered from at least 65% of the samples, analyzing each of the tissue-types and ant species separately using the microbiome v1.16 R package (68). We constructed heatmaps to visualize variation in relative abundance of symbionts (i.e., *Wolbachia*, *Spiroplasma* and *Mesoplasma*) in eggs and queen and gyne body samples using the packages phyloseq and NeatMap v0.3.6.2 (69). We also constructed bar plots using ggplot to visualize phylum-level bacterial microbiota composition for each tissue sampled in each ant colony.

Beta-diversity, or dissimilarity between samples, was assessed using Bray-Curtis dissimilarity and weighted Unifrac distances and evaluated using analysis of similarity (ANOSIM) and permutational multivariate analysis of variance (PERMANOVA, permutation number = 999) in vegan. Pairwise comparisons were performed with Bray-Curtis dissimilarity and weighted Unifrac distances and subjected to Bonferroni adjustment (package RVAideMemoire v0.9.81, function pairwise.perm.manova) (70). Worker mesosoma and gut communities were not significantly different from those of queens or gynes and were excluded from further beta-diversity comparisons to focus on vertically transmitted bacteria. Differences in beta-diversity were visualized using the Principal Coordinate Analysis (PCoA) ordination method using the R package vegan. Similarity percentage analysis (SIMPER) based on Bray-Curtis dissimilarity distances implemented in the vegan package followed by non-parametric Kruskal-Wallis tests with FDR corrected p-values were used to identify OTUs responsible for statistically significant differences in bacterial communities across ant species, tissues, and colonies within species (function: simper.pretty v1.1 in R doi.org/10.5281/zenodo.4270481) (71). We then constructed a heatmap to visualize relative abundances of the OTUs which significantly contributed to differences in beta-diversity of pellet bacterial microbiomes across ant species. For ecologically relevant clustering in this heatmap, the y-axis was ordered with Metric Multi-dimensional Scaling (MDS) using weighted Unifrac distances rather than by hierarchical cluster analysis which has no biological meaning in this case (69).

## Supplemental Material

**S1 Fig.**
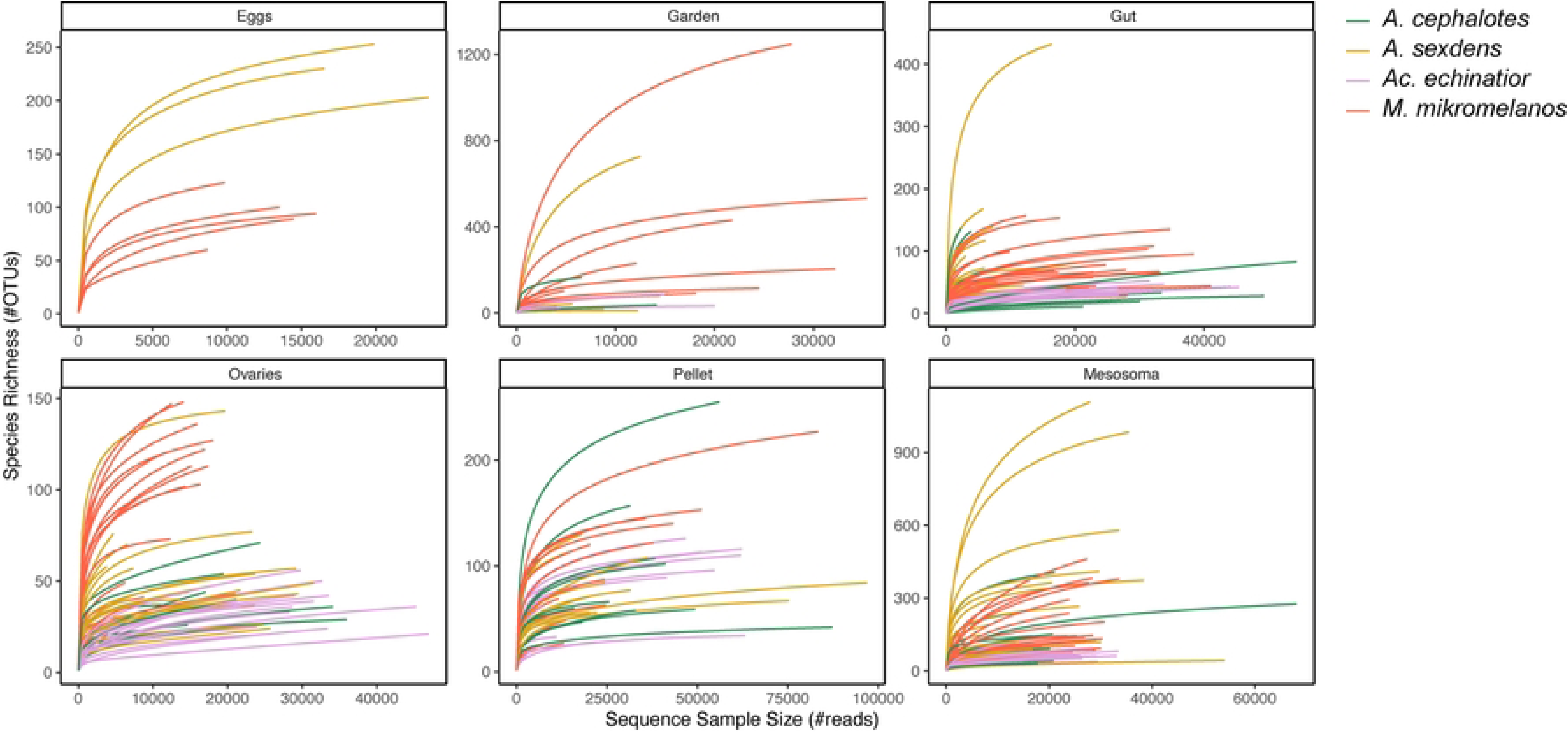
Rarefaction analyses of observed species richness as a function of sequencing depth for each tissue examined. Color indicates ant species. All samples in this study are represented, including those omitted from downstream analysis (e.g., workers and samples with <5000 reads).

**S2 Fig.**
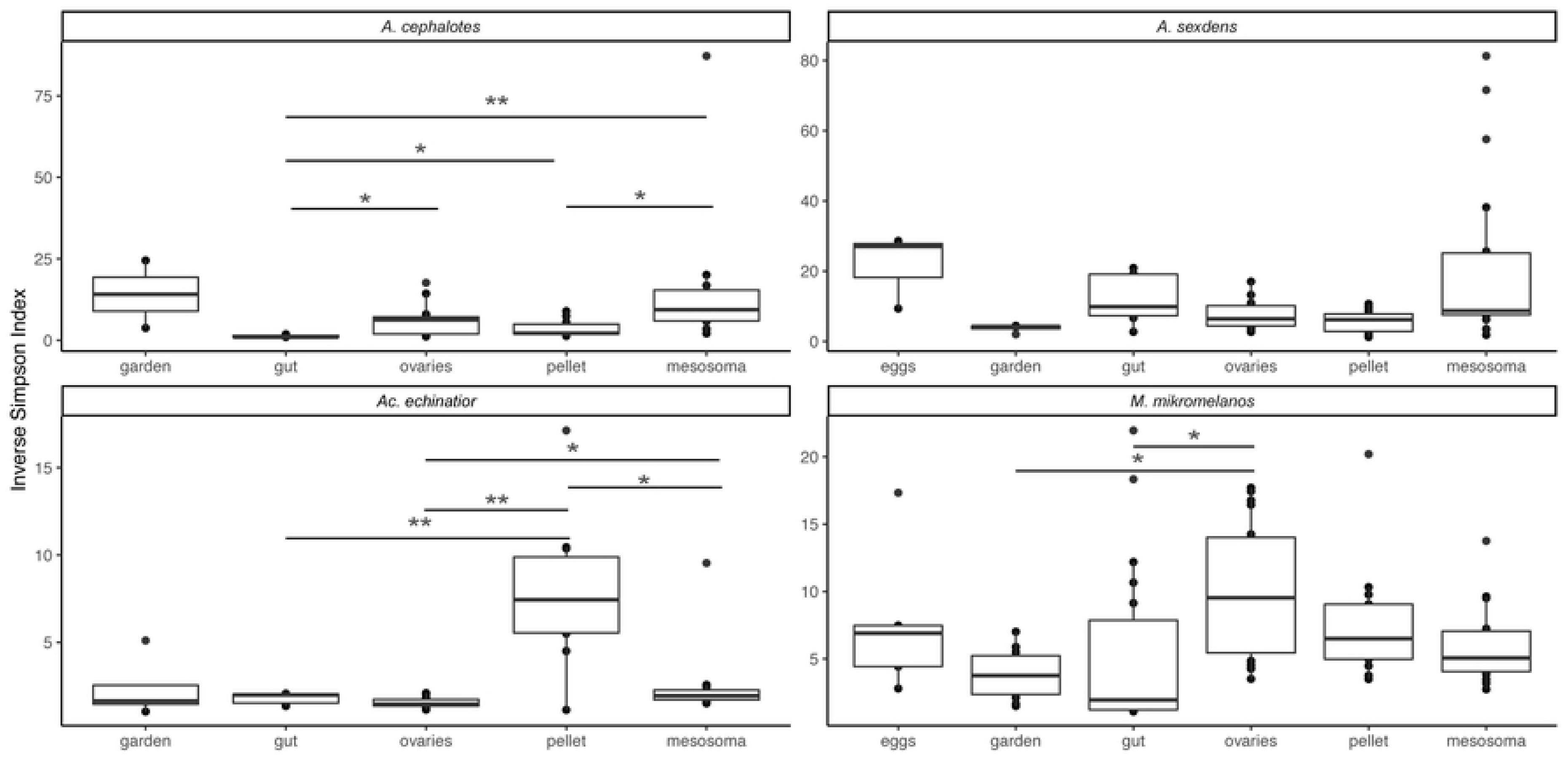
Comparison of alpha-diversity using the Inverse Simpson index of bacterial communities associated with queens across sample types within four ant species. Bars above plots indicate significance values for non-parametric Kruskal-Wallis and post hoc Wilcoxon rank sum tests for differences between species within each tissue type (* p ≤ 0.05, ∗∗ p ≤ 0.01). Note y-axis scale differs across plots. Sample sizes: Eggs (Mm: n = 5, As: n = 3), Ovaries (Mm: n = 18, Ae: n = 13, Ac: n = 13, As: n = 16), Garden (Mm: n = 10, Ae: n = 4, Ac: n = 2, As: n = 4), Pellet (Mm: n = 13, Ae: n = 11, Ac: n = 10, As: n = 13), Gut (Mm: n = 20, Ae: n = 13, Ac: n = 6, As: n = 9), Mesosoma (Mm: n = 20, Ae: n = 12, Ac: n = 10, As: n = 19).

**S3 Fig.**
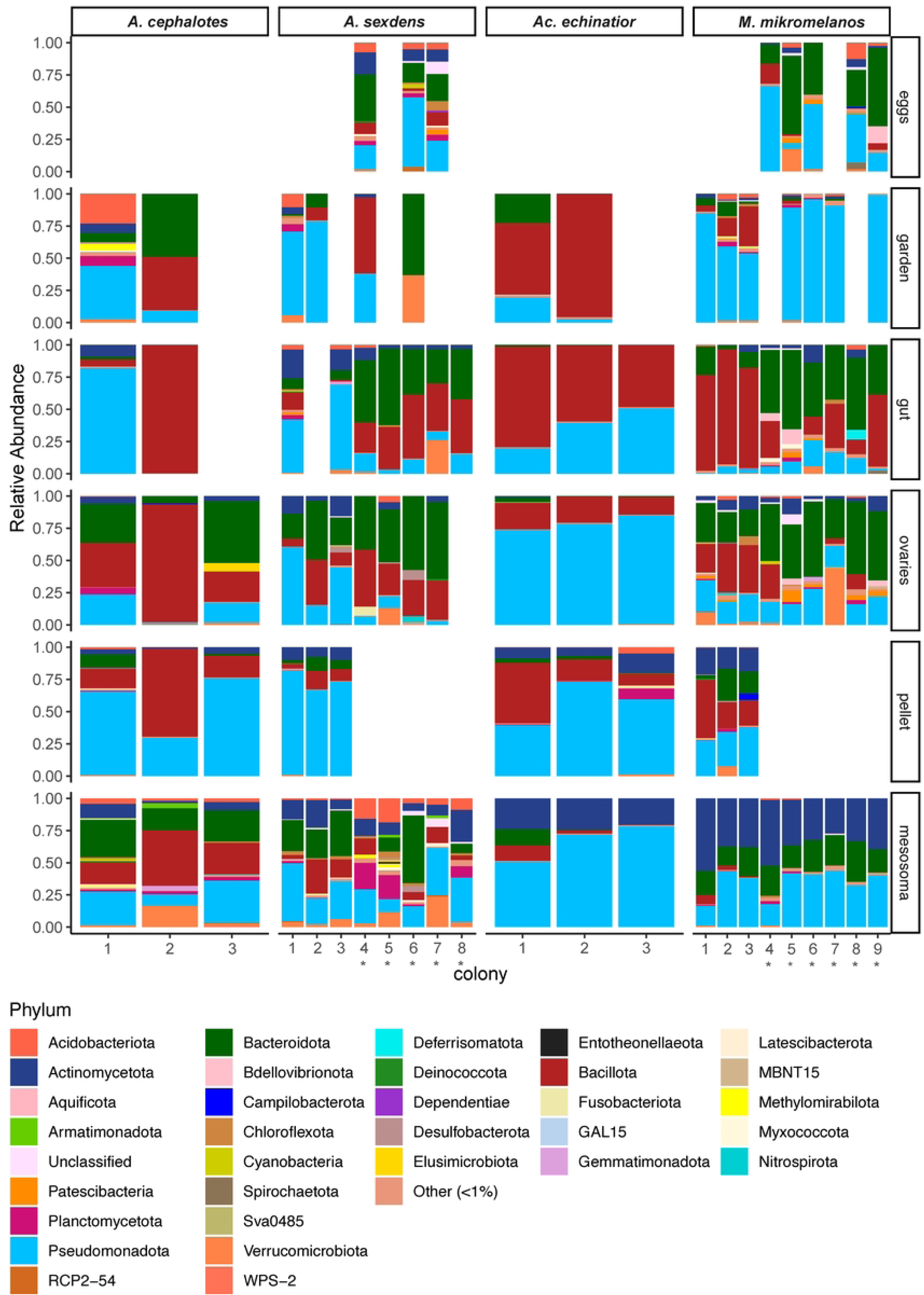
Bacterial microbiomes of gyne and queen tissues in four fungus-farming ant species characterized by phyla. Samples are pooled by colony and species identifiers (As, Ac, Ae and Mm) on colony IDs are omitted for simplicity. Samples derived from young queens rather than gynes are noted with an asterisk (*). Phyla which comprised <1% of a community in each sample were binned into a separate category.

**S1 Table. Metadata for each sample (n=301).**

Samples highlighted in yellow were excluded from beta-diversity analyses since the total reads was <5,000 for those samples.

**S2 Table. Closest NCBI RefSeq relatives to sequences from top 50 OTUs.**

Top OTUs were chosen from rarefied dataset.

**S3 Table. Taxonomic assignment and absolute abundance of bacterial OTUs observed.**

Samples highlighted in yellow are blank controls.

**S4 Table. OTUs (at 97% sequence similarity) inferred to belong to the core microbiome within sample types examined for four fungus-farming ant species.**

Core microbiota are defined as taxa present in at least 65% of samples within a group. *Mesoplasma* and gut core analyses are restricted to queen samples.

**S5 Table. Alpha diversity statistics using the Inverse Simpson measure.**

Statistically significant values (p<0.05) are bolded.

**S6 Table. Permutational multivariate analysis of variance (PERMANOVA) using weighted Unifrac distances comparing bacterial communities.** A) Comparison of pellet microbiota between ant species and between colonies within species. B) Comparison of garden, eggs and queen tissues between ant species. C) Comparison of pellet/garden samples and egg/ovaries samples within ant species. Statistically significant values (p<0.05) are bolded.

**S7 Table. Permutational multivariate analysis of variance (PERMANOVA) using Bray-Curtis dissimilarity distances comparing bacterial communities.** A) Comparison of pellet microbiota between ant species and between colonies within species. B) Comparison of garden, eggs and queen tissues between ant species. C) Comparison of pellet/garden samples and egg/ovaries samples within ant species. Statistically significant values (p<0.05) are bolded.

**S8 Table. SIMPER analysis indicating significant contributions of individual OTUs to observed differences in beta-diversity in pellets between ant species.** Statistically significant values (p<0.05) are bolded.

## Acknowledgments

We are grateful for permit and facilities support from the Smithsonian Tropical Research Institute (STRI) and we thank Zakee Sabree and Alison Bennett for valuable feedback on an earlier version of this manuscript. Ideas and insights were partly gained as an outcome of the 2019 Tropical Behavioral Ecology and Evolution course (OSU Global Education, Office of International Affairs; EEOB5797). This work was supported by The Ohio State University and funded by the National Science Foundation (RMMA: DEB SBS-2146104 and IOS-BS & PSS-2127521; https://www.nsf.gov/), the Brazilian Federal Agency for Support and Evaluation of Graduate Education (PK: CAPES-PrInt, process number 88887.310463/2018–00; #88887.468939/2019-00 and #88887.571230/2020–00) and Fundação de Amparo à Pesquisa do Estado de São Paulo (PK: #2019/22329-0 and #2022/14456-5). Funding institutions named did not play a role in the study design, data collection and analysis, decision to publish, or preparation of the manuscript. We also thank # anonymous reviewers for their valuable feedback on the manuscript.

## References

1. Bronstein JL. Mutualism. Oxford, UK: Oxford University Press; 2015.

2. Foster KR, Wenseleers T. A general model for the evolution of mutualisms. J Evol Biol. 2006;19(4):1283–93.

3. Harcombe WR, Chacón JM, Adamowicz EM, Chubiz LM, Marx CJ. Evolution of bidirectional costly mutualism from byproduct consumption. Proc Natl Acad Sci USA. 2018;115(47):12000–4.

4. Kikuchi Y, Hosokawa T, Fukatsu T. An ancient but promiscuous host-symbiont association between *Burkholderia* gut symbionts and their heteropteran hosts. ISME J. 2011;5:446–60.

5. Li YH, Ahmed MZ, Li SJ, Lv N, Shi PQ, Chen XS, et al. Plant-mediated horizontal transmission of *Rickettsia* endosymbiont between different whitefly species. FEMS Microbiol Ecol. 2017;93:138.

6. Visick KL, Stabb E V., Ruby EG. A lasting symbiosis: how *Vibrio fischeri* finds a squid partner and persists within its natural host. Nat Rev Microbiol. 2021;19(10):654–65.

7. Leftwich PT, Chapman T, McDonald J, Koskella B, Julian Marchesi P, Edgington MP. Transmission efficiency drives host-microbe associations. Proceedings of the Royal Society B: Biological Sciences. 2020;287(1934).

8. Bright M, Bulgheresi S. A complex journey: Transmission of microbial symbionts. Nature Reviews Microbiology. 2010;8:218–30.

9. Onchuru TO, Javier Martinez A, Ingham CS, Kaltenpoth M. Transmission of mutualistic bacteria in social and gregarious insects. Curr Opin Insect Sci. 2018;28:50–8.

10. Zhukova M, Sapountzis P, Schiøtt M, Boomsma JJ. Diversity and transmission of gut bacteria in *Atta* and *Acromyrmex* leaf-cutting ants during development. Front Microbiol. 2017;8(1942).

11. Weber NA. Fungus-growing ants. Science. 1966;153(3736):587–604.

12. De Fine Licht HH, Boomsma JJ. Forage collection, substrate preparation, and diet composition in fungus-growing ants. Ecol Entomol. 2010;35(3):259–69.

13. Solomon SE, Rabeling C, Sosa-Calvo J, Lopes CT, Rodrigues A, Vasconcelos HL, et al. The molecular phylogenetics of *Trachymyrmex* Forel ants and their fungal cultivars provide insights into the origin and coevolutionary history of ‘higher-attine’ ant agriculture. Syst Entomol. 2019;44(4):939–56.

14. Suen G, Scott JJ, Aylward FO, Adams SM, Tringe SG. An insect herbivore microbiome with high plant biomass-degrading capacity. PLoS Genet. 2010;6(9):1001129.

15. Aylward FO, Burnum KE, Scott JJ, Suen G, Tringe SG, Adams SM, et al. Metagenomic and metaproteomic insights into bacterial communities in leaf-cutter ant fungus gardens. ISME J. 2012;6:1688–1701.

16. Aylward FO, Burnum-Johnson KE, Tringe SG, Teiling C, Tremmel DM, Moeller JA, et al. *Leucoagaricus gongylophorus* produces diverse enzymes for the degradation of recalcitrant plant polymers in leaf-cutter ant fungus gardens. Appl Environ Microbiol. 2013;79(12).

17. Moreira-Soto RD, Sanchez E, Currie CR, Pinto-Tomás AA. Ultrastructural and microbial analyses of cellulose degradation in leaf-cutter ant colonies. Microbiology. 2017;163(11):1578–89.

18. Huang EL, Aylward FO, Kim YM, Webb-Robertson BJM, Nicora CD, Hu Z, et al. The fungus gardens of leaf-cutter ants undergo a distinct physiological transition during biomass degradation. Environ Microbiol Rep. 2014;6(4):389–95.

19. Shik JZ, Kooij PW, Donoso DA, Santos JC, Gomez EB, Franco M, et al. Nutritional niches reveal fundamental domestication trade-offs in fungus-farming ants. Nat Ecol Evol. 2021;5(1):122–34.

20. Scott JJ, Budsberg KJ, Suen G, Wixon DL, Balser TC, Currie CR. Microbial community structure of leaf-cutter ant fungus gardens and refuse dumps. PLoS One. 2010;5(3).

21. Pinto-Tomás AA, Anderson MA, Suen G, Stevenson DM, Chu FST, Cleland WW, et al. Symbiotic nitrogen fixation in the fungus gardens of leaf-cutter ants. Science. 2009;326(5956):1120–3.

22. Meirelles LA, Mcfrederick QS, Rodrigues A, Mantovani JD, de Melo Rodovalho C, Ferreira H, et al. Bacterial microbiomes from vertically transmitted fungal inocula of the leaf-cutting ant *Atta texana*. Environ Microbiol Rep. 2016;8:630–40.

23. Kellner K, Ishak HD, Linksvayer TA, Mueller UG. Bacterial community composition and diversity in an ancestral ant fungus symbiosis. FEMS Microbiol Ecol. 2015;91(7):fiv073.

24. Ronque MUV, Lyra ML, Migliorini GH, Bacci M, Oliveira PS. Symbiotic bacterial communities in rainforest fungus-farming ants: evidence for species and colony specificity. Sci Rep. 2020;10(1):10172.

25. Ishak HD, Miller JL, Sen R, Dowd SE, Meyer E, Mueller UG. Microbiomes of ant castes implicate new microbial roles in the fungus-growing ant *Trachymyrmex septentrionalis*. Sci Rep. 2011;1:204.

26. Haeder S, Wirth R, Herz H, Spiteller D. Candicidin-producing *Streptomyces* support leaf-cutting ants to protect their fungus garden against the pathogenic fungus *Escovopsis*. Proc Natl Acad Sci USA. 2009;106(12):4742–6.

27. Barke J, Seipke RF, Grüschow S, Heavens D, Drou N, Bibb MJ, et al. A mixed community of actinomycetes produce multiple antibiotics for the fungus farming ant *Acromyrmex octospinosus*. BMC Biol. 2010;8(109).

28. Andersen SB, Hansen LH, Sapountzis P, Sørensen SJ, Boomsma JJ. Specificity and stability of the *Acromyrmex*-*Pseudonocardia* symbiosis. Mol Ecol. 2013;22(16):4307–21.

29. Mueller UG, Ishak H, Lee JC, Sen R, Gutell RR. Placement of attine ant-associated *Pseudonocardia* in a global *Pseudonocardia* phylogeny (Pseudonocardiaceae, Actinomycetales): a test of two symbiont-association models. Antonie Van Leeuwenhoek. 2010;98(2):195–212.

30. Mattoso TC, Moreira DDO, Samuels RI. Symbiotic bacteria on the cuticle of the leaf-cutting ant *Acromyrmex subterraneus subterraneus* protect workers from attack by entomopathogenic fungi. Biol Lett. 2012;8(3):461–4.

31. Sapountzis P, Nash DR, Schiøtt M, Boomsma JJ. The evolution of abdominal microbiomes in fungus-growing ants. Mol Ecol. 2019;28(4):879–99.

32. Yun JH, Roh SW, Whon TW, Jung MJ, Kim MS, Park DS, et al. Insect gut bacterial diversity determined by environmental habitat, diet, developmental stage, and phylogeny of host. Appl Environ Microbiol. 2014;80(17):5254–64

33. Colman DR, Toolson EC, Takacs-Vesbach CD. Do diet and taxonomy influence insect gut bacterial communities? Mol Ecol. 2012;21(20):5124–37.

34. Sapountzis P, Zhukova M, Hansen LH, Sørensen SJ, Schiøtt M, Boomsma JJ. *Acromyrmex* leaf-cutting ants have simple gut microbiota with nitrogen-fixing potential. Appl Environ Microbiol. 2015;81(16):5527–37.

35. Frost CL, Fernández Marín H, Smith JE, Hughes WOH. Multiple gains and losses of *Wolbachia* symbionts across a tribe of fungus-growing ants. Mol Ecol. 2010;19(18):4077– 85.

36. Andersen SB, Boye M, Nash DR, Boomsma JJ. Dynamic *Wolbachia* prevalence in *Acromyrmex* leaf-cutting ants: Potential for a nutritional symbiosis. J Evol Biol. 2012;25(7):1340–50.

37. Liberti J, Sapountzis P, Hansen LH, Sørensen SJ, Adams RMM, Boomsma JJ. Bacterial symbiont sharing in *Megalomyrmex* social parasites and their fungus-growing ant hosts. Mol Ecol. 2015;24(12):3151–69.

38. Nygaard S, Hu H, Li C, Schiøtt M, Chen Z, Yang Z, et al. Reciprocal genomic evolution in the ant-fungus agricultural symbiosis. Nat Commun. 2016;7.

39. Sapountzis P, Zhukova M, Shik JZ, Schiott M, Boomsma JJ. Reconstructing the functions of endosymbiotic mollicutes in fungus-growing ants. Elife. 2018;7.

40. Zhukova M, Sapountzis P, Schiøtt M, Boomsma JJ. Phylogenomic analysis and metabolic role reconstruction of mutualistic Rhizobiales hindgut symbionts of *Acromyrmex* leaf-cutting ants. FEMS Microbiol Ecol. 2022;98:1–13.

41. Linnæus C. Systema naturae per regna tria naturae, secundum classes, ordines, genera, species, cum characteribus, differentiis, synonymis, locis. 10^th^ ed. Stockholm: Laurentii Salvii; 1758.

42. Forel A. Formicidae. In: Biologica Central-Americana Hymenoptera. London: R. H. Porter; 1899. pp. 25–56.

43. Cardenas CR, Luo AR, Jones TH, Schultz TR, Adams RMM. Using an integrative taxonomic approach to delimit a sibling species, Mycetomoellerius mikromelanos sp. nov. (Formicidae: Attini: Attina). PeerJ. 2021;9:e11622.

44. Chang PT, Rao K, Longo LO, Lawton ES, Scherer G, Van Arnam EB. Thiopeptide defense by an ant’s bacterial symbiont. J Nat Prod. 2020;83(3):725–9.

45. Cafaro MJ, Poulsen M, Little AEF, Price SL, Gerardo NM, Wong B, et al. Specificity in the symbiotic association between fungus-growing ants and protective *Pseudonocardia* bacteria. Proc. R. Soc. B. 2011;278(1713):1814–22.

46. Martiarena MJS, Deveau A, Montoya QV, Flórez L V., Rodrigues A. The hyphosphere of leaf-cutting ant cultivars is enriched with helper bacteria. Microb Ecol. 2023;86(3):1773– 88.

47. Chuah LO, Yap KP, Syuhada AKS, Thong KL, Ahmad R, Liong MT, et al. *Floricoccus tropicus* gen. nov., sp. nov. and *Floricoccus penangensis* sp. nov. isolated from fresh flowers of durian tree and hibiscus. Int J Syst Evol Microbiol. 2017;67(12):4979–85.

48. Russell JA, Funaro CF, Giraldo YM, Goldman-Huertas B, Suh D, Kronauer DJC, et al. A veritable menagerie of heritable bacteria from ants, butterflies, and beyond: broad molecular surveys and a systematic review. PLoS One. 2012;7(12):e51027.

49. Bringhurst B, Greenwold M, Kellner K, Seal JN. Symbiosis, dysbiosis and the impact of horizontal exchange on bacterial microbiomes in higher fungus-gardening ants. Sci Rep. 2024;14(1):3231.

50. de Oliveira Aquino Zani R, Ferro M, Bacci Jr M, Aquino Zani or Maurício Bacci Jr O. Three phylogenetically distinct and culturable diazotrophs are perennial symbionts of leaf-cutting ants. Ecology and Evolution. 2021;11:17686–99.

51. Rose C, Lund MB, Søgård AM, Busck MM, Bechsgaard JS, Schramm A, et al. Social transmission of bacterial symbionts homogenizes the microbiome within and across generations of group-living spiders. ISME Communications. 2023;3(1).

52. Ramalho MO, Moreau CS. Untangling the complex interactions between turtle ants and their microbial partners. Anim Microbiome. 2023;5(1).

53. Valencia-Giraldo SM, Niño-Castro A, López-Peña A, Trejos-Vidal D, Correa-Bueno O, Montoya-Lerma J. Immunity and survival response of *Atta cephalotes* (Hymenoptera: Myrmicinae) workers to *Metarhizium anisopliae* infection: Potential role of their associated microbiota. PLoS One. 2021;16(2).

54. Barcoto MO, Carlos-Shanley C, Fan H, Ferro M, Nagamoto S, Bacci M, et al. Fungus-growing insects host a distinctive microbiota apparently adapted to the fungiculture environment. Sci Rep. 2020;10:12384.

55. Bringhurst B, Allert M, Greenwold M, Kellner K, Seal JN. Environments and hosts structure the bacterial microbiomes of fungus-gardening ants and their symbiotic fungus gardens. Microb Ecol. 2022;86:1374–92.

56. Branstetter MG, Ješovnik A, Sosa-Calvo J, Lloyd MW, Faircloth BC, Brady SG, et al. Dry habitats were crucibles of domestication in the evolution of agriculture in ants. Proc. R. Soc. B. 2017;284(1852):20170095.

57. Mueller UG, Kardish MR, Ishak HD, Wright AM, Solomon SE, Bruschi SM, et al. Phylogenetic patterns of ant-fungus associations indicate that farming strategies, not only a superior fungal cultivar, explain the ecological success of leafcutter ants. Mol Ecol. 2018;27(10):2414–34.

58. Mueller UG, Ishak HD, Bruschi SM, Smith CC, Herman JJ, Solomon SE, et al. Biogeography of mutualistic fungi cultivated by leafcutter ants. Mol Ecol. 2017;26(24):6921–6937.

59. González CT, Saltonstall K, Fernández-Marín H. Garden microbiomes of *Apterostigma dentigerum* and *Apterostigma pilosum* fungus-growing ants (Hymenoptera: Formicidae). Journal of Microbiology. 2019;57(10):842–51.

60. Khadempour L, Fan H, Keefover-Ring K, Carlos-Shanley C, Nagamoto NS, Dam MA, et al. Metagenomics reveals diet-specific specialization of bacterial communities in fungus gardens of grass- and dicot-cutter ants. Front Microbiol. 2020;11:2227.

61. Schiøtt M, De Fine Licht HH, Lange L, Boomsma JJ. Towards a molecular understanding of symbiont function: Identification of a fungal gene for the degradation of xylan in the fungus gardens of leaf-cutting ants. BMC Microbiol. 2008;8(1):1–10.

62. Kozich JJ, Westcott SL, Baxter NT, Highlander SK, Schloss PD. Development of a dual-index sequencing strategy and curation pipeline for analyzing amplicon sequence data on the MiSeq Illumina sequencing platform. Appl Environ Microbiol. 2013;79(17):5112–20.

63. Schloss PD, Westcott SL, Ryabin T, Hall JR, Hartmann M, Hollister EB, et al. Introducing mothur: Open-source, platform-independent, community-supported software for describing and comparing microbial communities. Appl Environ Microbiol. 2009;75(23):7537–41.

64. Oksanen FJ, Blanchet G, Friendly M, Kindt R, Legendre P, McGlinn D, et al. Vegan: Community Ecology Package. R version 2.5–7. 2020. Available from: https://CRAN.R-project.org/package=vegan

65. McMurdie PJ, Holmes S. Phyloseq: An R Package for Reproducible Interactive Analysis and Graphics of Microbiome Census Data. PLoS One. 2013;8(4).

66. Kandlikar GS, Gold ZJ, Cowen MC, Meyer RS, Freise AC, Kraft NJB, et al. ranacapa: An R package and Shiny web app to explore environmental DNA data with exploratory statistics and interactive visualizations. 2018; Available from: 10.12688/f1000research.16680.1

67. Lahti L, Shetty S, Salojarvi J. Tools for microbiome analysis in R. Microbiome package version 1.17.42. 2017; Available from: https://github.com/microbiome/microbiome

68. Rajaram S, Oono Y. NeatMap - non-clustering heat map alternatives in R. BMC Bioinformatics. 2010;11(1):45.

69. Hervé M. RVAideMemoire: testing and plotting procedures for biostatistics. R package version 0.9-70. 2018; Available from: https://cran.r-project.org/web/packages/RVAideMemoire/index.html

70. asteinberger9. asteinberger9/seq_scripts v1.1. 2020; Available from: https://zenodo.org/record/4270481

